# Proteomic characterisation of the early rheumatoid arthritis-cardiovascular disease multimorbid axis

**DOI:** 10.1101/2025.02.17.638761

**Authors:** Rudresh R Shukla, Amr Mohammed, Nicholas Black, Bara Erhayiem, Graham Fent, Christopher A Miller, Sven Plein, Darren Plant, Maya H Buch

## Abstract

**Background:** Inflammation contributes to the increased risk of cardiovascular disease (CVD) observed in people with rheumatoid arthritis (RA), with increased prevalence observed from time of diagnosis. Subclinical vascular and myocardial abnormalities can be detected with cardiovascular magnetic resonance (CMR) imaging in otherwise low risk individuals. Identifying associated circulating markers could enable diagnostics and implicate biological pathways of RA-CVD. The study’s objective was to identify blood-based proteins associated with CMR measures of vascular and myocardial abnormality and associated inflammatory and/or cardio-metabolic pathways in a new-onset RA cohort.

**Methods:** Serum samples (baseline/pre-treatment, N = 75 and year 1, N = 71) from CADERA (Coronary Artery Disease Evaluation in Rheumatoid Arthritis) participants, a subgroup of a randomised controlled trial, who underwent CMR, were used to measure 334 proteins across 4 pre-defined Olink panels (Inflammation, Cardiovascular-II, Cardiovascular-III, Cardiometabolic). Bayesian mixed effects regression analyses, unsupervised hierarchical clustering and protein network analyses were applied. Normalised protein expression from 334 Olink proteins were used as exposures in the regression analyses. CMR measures of vascular stiffness (aortic distensibility and stiffness index) and myocardial tissue characteristics (native T1, myocardial extracellular volume and late gadolinium enhancement) were each used as individual outcomes in regression analyses. Associations were considered significant at 95% threshold for credible intervals. An expanded physical protein-protein interaction (PPI) network created using the significant (seed) proteins was subjected to topological and enrichment analysis to identify enriched biological pathways.

**Results:** 54/334 proteins were associated significantly with CMR measures at baseline (7 - vascular; 48 - myocardial tissue characteristics; Coagulation factor 11 with both). Two proteins, TNFSF13B (B-cell activating factor) and CRTAC1, were associated with CMR measures at baseline, were sensitive to change over time and co-varied with changes in CMR measure. Topological analysis of expanded PPI network revealed GRB2 as most connected (degree score=64; closeness score=118.83) and four key signalling pathways, including JAK-STAT and EGFR tyrosine kinase inhibitor resistance, emerged as significant.

**Conclusions:** This first proteomic study of treatment-naïve early RA and subclinical cardiovascular pathology identifies proteins that could aid diagnostic test development and implicates signalling pathways in the RA-cardiovascular axis to inform on our understanding of the basis of RA-CVD.

## Introduction

Rheumatoid arthritis (RA) is associated with higher mortality compared to the general population, primarily due to an increased risk of cardiovascular disease (CVD) secondary to accelerated atherosclerosis and heart failure (HF) ^1–4^. Whilst earlier diagnosis of RA and effective targeted therapies have improved cardiovascular (CV) outcomes^5^, specific risk models and stratification to aid early detection and intervention are lacking. An appreciation of shared inflammatory pathways across RA/inflammatory diseases and atherosclerotic CVD (ASCVD) such as interleukin (IL) −1 and IL-6 signalling has emerged ^6^ but a more refined understanding to inform precision medicine use of therapies in RA is still needed^7^.

Cardiovascular magnetic resonance imaging (CMR) is highly effective in characterising myocardial tissue changes, detailing vasculature and myocardial function, and quantifying changes in these measurements ^8^. CMR studies have reported subclinical vascular and myocardial abnormalities in individuals with very early RA (ERA) ^9^ and established RA ^10^. This allows detection of early processes of ASCVD and HF that are associated with adverse outcomes in a range of patient populations ^11–13^ and may provide a basis for investigating underlying biological pathways. However, screening of unselected patients using such imaging techniques is not appropriate.

High throughput proteomic techniques offer considerable opportunities to understand disease complexity^14,15^. Recent studies focussing on RA patients and HF, using inflammation or CVD-related proteins, have provided initial insights into the association between chronic inflammation and HF risk ^16,17^. There remains limited insight into the pathways underlying specific CV processes early in RA to inform effective diagnostics, progress mechanistic understanding and enable optimal therapeutic strategies.

Within a treatment-naïve, new-onset ERA randomised controlled trial cohort, this study aimed to identify blood-based protein biomarkers that associate with CMR measures of vascular stiffness (VS) and abnormal myocardial tissue characteristics, evaluate whether these biomarkers were sensitive to change in CMR measures following treatment for RA and the predominant inflammatory and/or cardio-metabolic pathways implicated.

## Methods

### Study design

A biomarker study of the ‘CADERA’ (Coronary Artery Disease Evaluation in Rheumatoid Arthritis) cohort ^9^, a subgroup of the phase IV ERA trial (‘VEDERA’)^18^ randomised to etanercept and methotrexate (ETN+MTX) or MTX treat-2-target strategy.

### Patient population

‘VEDERA’ participants had a diagnosis of treatment-naïve, ERA. The CADERA study subgroup participants did not have a history of CVD and had low cardiovascular risk scores with a maximum of one traditional risk factor (TRF) permitted [excluding diabetes mellitus (DM)].

### Assessments

Individuals underwent CMR to detect subclinical CV abnormalities at baseline (pre-treatment) and following randomised treatment at year 1. Primary outcomes from both VEDERA^18^ and CADERA^9^ have been published.

### Cardiovascular magnetic resonance imaging measures and serum cardiac biomarker testing

All scans were performed on the same 3 Tesla system and the protocol has been previously reported^9^. Outcomes of interest for this study were CMR measures of VS [aortic distensibility (AD), aortic stiffness index (AoSI)] and myocardial tissue characteristics [native T1, myocardial extracellular volume (ECV) and late gadolinium enhancement (LGE)]. In addition, high sensitivity troponin I (hsTnI) and N-terminal Pro-B type natriuretic peptide (NTproBNP) levels were measured at baseline and year 1.

### Olink Proteomics

Serum samples collected at time of CMR (baseline, n=75 and Year 1, n=71) were sent to an Olink accredited lab (University of Leeds; www.olink.com) and analysed across 4 pre-defined Target-96 panels – Cardiometabolic (CM), Cardiovascular-II (CV2), Cardiovascular-III (CV3) and Inflammation (INF) ^19^ using proximity extension assay technique with extensive built-in intra and inter-plate controls.

### Statistical analysis

Data from Olink are reported back as normalised protein expression (NPX) values on a log2 scale. Proteins with more than 40% of samples that were below the lower limit of detection (LLOD) were excluded from analyses ^20^. Bayesian mixed effects regression (Rstanarm package, v 2.21.4) ^21^ was used to identify proteins associated with individual CMR measures [AD, AoSI, native T1, ECV are continuous; LGE is categorical (presence/absence)] at baseline. For each protein, in addition to NPX, age, gender, systolic blood pressure and number of pack-years smoked were included as fixed effects and participant ID was included as a random effect. Model performance was assessed using the loo package (v 2.6.0). Including body mass index did not improve the predictive performance of the model. To address dependency on time-point, an interaction term between NPX and time-point (categorical) was included. To identify proteins sensitive to change over time, a separate analysis was performed using NPX as outcome and time-point as a fixed effect in addition to other fixed and random effects listed above. Results from the Bayesian regression analyses are presented as posterior estimates (PE) with 95% credible intervals (CrI). Results were defined as significant if the credible intervals did not include zero. Significant results with a positive posterior estimate suggest a positive association of the protein level with outcome (meaning, increase in protein levels with an increase in CMR measure) and vice versa, whilst a negative posterior estimate suggests an inverse association (meaning, increase in protein levels with a decrease in CMR measure or vice versa). All statistical analyses were conducted on R (v4.2.2).

### Physical protein-protein interaction network

Baseline proteins that were associated significantly with baseline CMR measures were expanded using the Cytoscape StringDB plugin ^22^ limiting to only a physical subnetwork (excluding co-expression or co-occurrence, confidence cutoff = 0.4). K-means limited to a minimum of 20 proteins within each cluster was used to identify potential sub-clusters within the expanded network. Cytoscape was used to assess structural properties of the overall expanded network and Cytohubba plugin was used for topological analysis to identify the top overall and within-cluster proteins by degree, betweenness and closeness^23^. The overall PPI network and sub-clusters were subjected to Gene ontology (GO) and Kyoto Encyclopedia of Genes and Genomes (KEGG) pathway analysis using ShinyGO ^24^.

## Results

Baseline demographic data have been published previously^9^ and are summarised in table 1. The median age was 51 years [inter-quartile range (IQR) 40 – 61] with 52/75 (69%) females, 40/75 (53%) current or ex-smokers and median body mass index (BMI) 26.4kg/m2 (IQR 23.3-28). Participants had a median RA symptom duration of 21.9 weeks (IQR 14.4-32.7) and, as per parent trial recruitment criteria, active RA disease [median disease activity score using erythrocyte sedimentation rate (DAS)28-ESR 5.51(4.88-6.40) and C-reactive protein (CRP) 8.38mg/L(2.3-20.48)] ^18^. In line with study eligibility, prevalence of TRF for CVD was low, 10% had a history of hypertension (HTN), hypercholesterolaemia or family history of ischaemic heart disease.

**Table 1:** Baseline clinical characteristics of participants. Values shown are Q50 (Q25-Q75) or percentages. BMI = body mass index, DAS28-ESR = Disease activity score (28-joints) with erythrocyte sedimentation rate, SDAI = Simplified Disease Activity Index, ESR = erythrocyte sedimentation rate, CRP = C-reactive protein, HAQ = Health assessment questionnaire score, NTproBNP = N-terminal fragment of B-type natriuretic peptide, Hs-TnI = high sensitivity troponin I. *Total PD score = sum of semiquantitative power Doppler scores from each joint scanned (dominant/most symptomatic metacarpophalangeal joints 1-5, wrist).

| Characteristic | Overall, N = 75 |
| --- | --- |
| Age (yrs) | 51 (40, 61) |
| Female | 52 / 75 (69%) |
| Current/ex-smokers | 40 / 75 (53%) |
| BMI (kg/m<sup>2</sup>) | 26 (23, 28) |
| Symptom duration (weeks) | 21.9 (14.4, 32.7) |
| Family history |  |
| Vascular disease | 16 / 75 (21%) |
| Ischaemic heart disease | 4 / 75 (5.3%) |
| Premature cardiovascular disease | 7 / 75 (9.3%) |
| Cardiovascular history |  |
| Hypertension | 6 / 75 (8.0%) |
| Hypercholesterolaemia | 2 / 75 (2.7%) |
| Ischaemic heart disease | 2 / 75 (2.7%) |
| Cerebrovascular disease | 0 / 75 (0%) |
| Peripheral vascular disease | 0 / 75 (0%) |
| Disease characteristics |  |
| seropositive | 67 / 75 (89%) |
| DAS28-ESR | 5.51 (4.88, 6.40) |
| SDAI | 29.18 (20.50, 39.56) |
| ESR | 32 (18, 54) |
| CRP | 8(2, 20) |
| Power Doppler on Ultrasound (total PD score > 0)* | 47 / 75 (63%) |
| HAQ | 1.26 (0.86, 1.49) |
| Cardiac biomarkers (clinically measured) |  |
| NTproBNP (pg/ml) | 72 (38, 119) |
| Hs-TnI (ng/L) | 2 (1, 3) |

### Proteins associated with CMR measures at baseline

In total, 334/368 proteins were detected across the 4 panels of which 318 were above the LLOD predetermined by Olink. Table S1 lists the percentage of samples below the LLOD for each protein.

At baseline, 54/318 proteins were associated significantly with CMR measures [7 with VS (AD and/or AoSI), 48 with myocardial tissue characteristics (native T1, ECV or LGE)] (figure 1, table S2). HB-EGF (CV2 panel) was associated with AD and AoSI vascular CMR measures (AD = PE −0.6, CrI −1.14 to −0.05; AoSI = PE 0.9, CrI 0.18 to 1.63). F11 (CM panel) was associated with both vascular and myocardial CMR measures (AoSI = PE −1.8, CrI −3.53 to −0.07; ECV = PE −0.027, CrI −0.054 to −0.004)].

**Figure 1:**
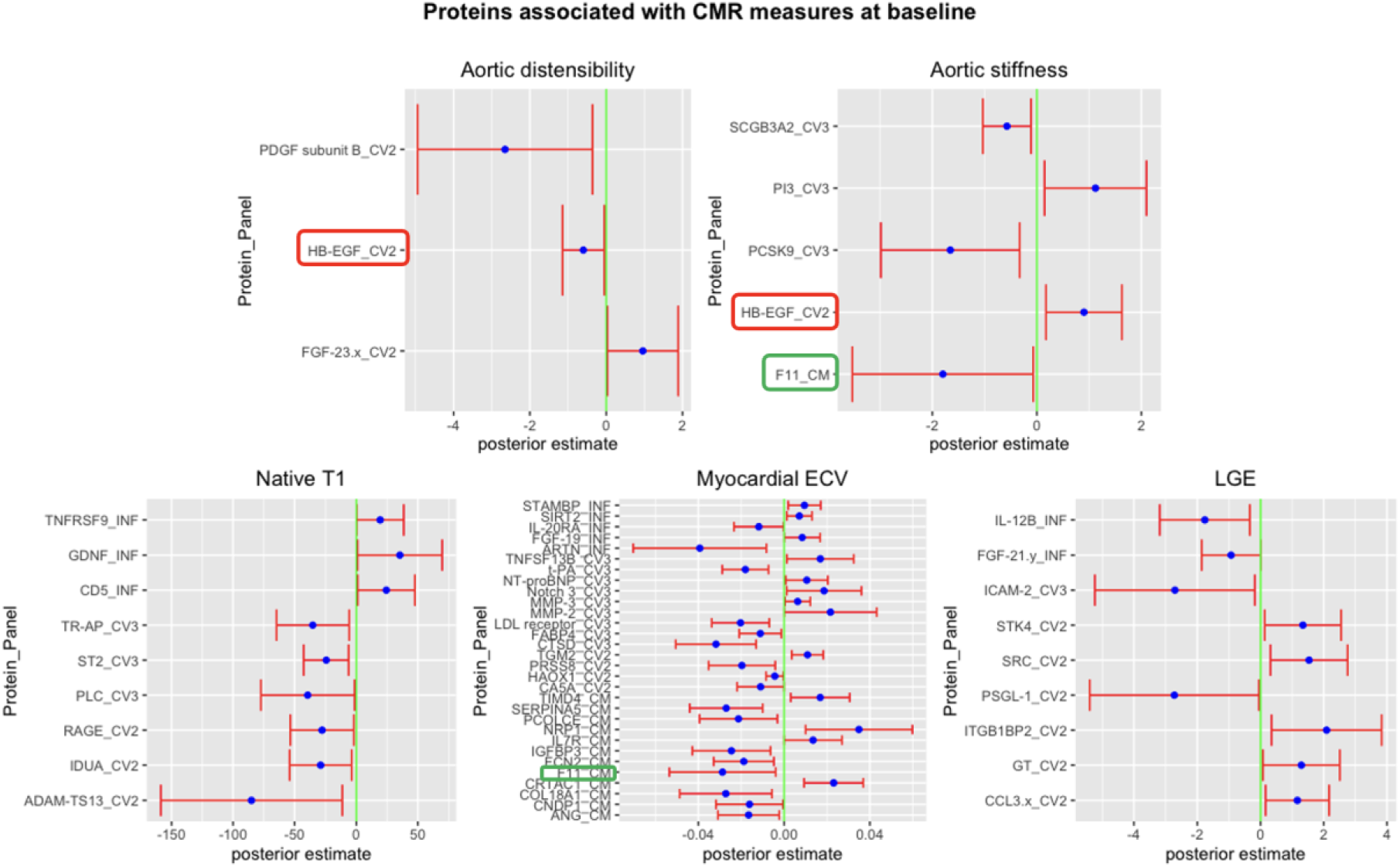
Proteins associated with different cardiovascular magnetic resonance (CMR) imaging measures at baseline. Y axis label (Protein_Panel) indicates protein followed by Olink panel name. 2 overlapping proteins across different CMR measures are F11 from the cardiometabolic panel (CM) and HB_EDF from cardiovascular II (CV2) panel. Posterior estimates (blue dot) with 95% credible intervals (red bar) reported. ECV = extracellular volume, LGE = late gadolinium enhancement.

Top proteins associated with VS, included PDGF subunit B [AD = PE −2.65, CrI −4.94 to −0.36] and F11 (AoSI = PE −1.8, CrI −3.53 to −0.07) and with myocardial tissue characteristics, ADAM-TS13 (native T1 = PE −85.05, CrI −158.7 to −11.4), ARTN (ECV = PE −0.04, CrI −0.07 to −0.008) and PSGL-1 (LGE = PE-2.73, −5.4 to −0.05). The 54 identified proteins were significantly correlated with each other (figure S1). Raw NPX values (pre-imputation) and clinic measured NT-proBNP levels were significantly and highly correlated (coefficient = 0.77, p < 0.005) thus providing orthogonal validation of NPX values.

### Proteins sensitive over time and with interval change in CMR measures

Of the 54 significant baseline proteins, 12 were sensitive to change over time and 8 had a statistically significant interaction term between NPX and time-point (baseline and year 1) (5 = VS; 3 = myocardial tissue characteristics, figure 2). Two proteins, TNFSF13B and CRTAC1 (neither from the INF panel), were sensitive to change over time and co-varied with changes in CMR measure (AD and ECV, respectively) between baseline and year 1 (figure 2, figure S2). Both SIRT2 and STAMBP were significantly and positively associated with ECV at baseline and demonstrated a negative association with ECV at year 1 (figure S3). All proteins that were sensitive to change over time are listed in table S3, whereas those with a statistically significant interaction term between NPX and time-point (baseline and year 1) are listed in table S4.

**Figure 2:**
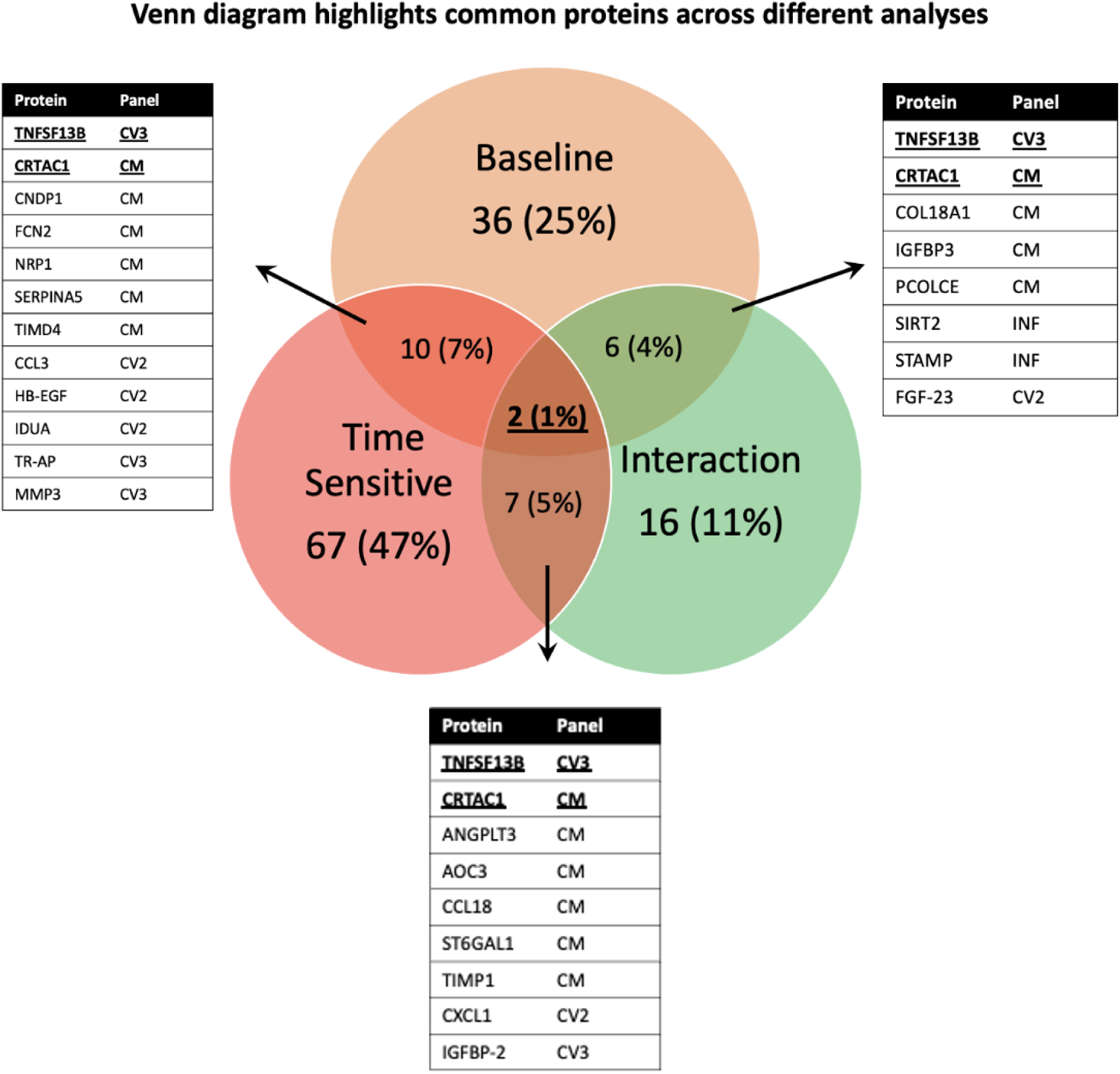
Venn diagram identifies common proteins across analyses – Baseline = proteins associated with CMR measures at baseline; Sensitive = Proteins sensitive to change over time; Interaction = Proteins with a difference in association with CMR measures at baseline and year 1; CRTAC1 and TNFSF13B are common across analyses.

### EGFR tyrosine kinase inhibitor resistance and JAK-STAT are key signalling pathways within the ERA-CVD expanded PPI network

The 53 baseline proteins (excluding NT-proBNP, figure 3A) were used to seed an overall physical expanded PPI network (figure 3B) that consisted of 203 nodes (proteins) and 1108 edges (interactions) (table S5). The overall network was highly connected (component score =15) and was likely to include functional modules or clusters of proteins (clustering coefficient = 0.49). GRB2 was the most connected protein overall (Table 2). Functional analysis revealed ‘peptidyl tyrosine phosphorylation’ as the top biological process and ‘EGFR tyrosine kinase inhibitor resistance’ [enrichment False Discovery Rate (FDR) = 7.3E-23] and ‘JAK-STAT signalling’ (FDR 3.9E-31) as the top enriched KEGG pathways for the expanded PPI network (figure S4, top). Expanded pathways contained several metabolic proteins such as FGF, PDGF, cytokines such as interleukin (IL)-2, −3, −10 family and IL-12/23 and growth factors (figure S4 middle and bottom, highlighted in red).

**Figure 3:**
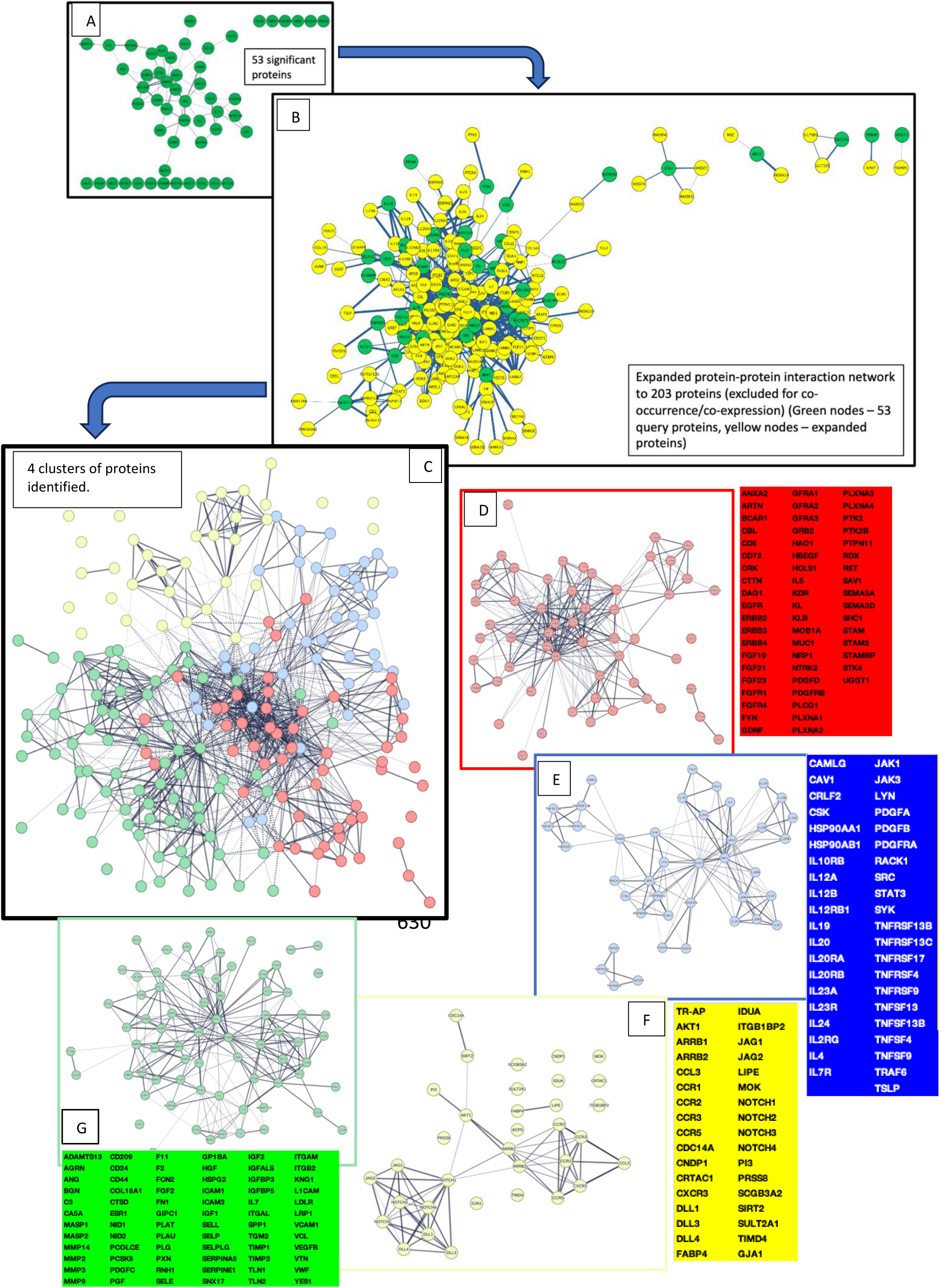
A – 53 proteins (green nodes) associated with CMR measures at baseline (excluding NTproBNP) used as seed proteins to create an expanded protein-protein interaction (PPI) network. B – A physical PPI network created (excluding co-occurrence and co-expression) (53 proteins expanded to 203 proteins). Green nodes indicate seed proteins from A. Yellow nodes are expanded proteins. C – K-means clustering applied to expanded PPI network and 4 clusters identified – red, blue, yellow, green D – Red cluster consists of 56 proteins listed in adjacent red table E – Blue cluster consists of 41 proteins listed in adjacent blue table F – Yellow cluster consists of 34 proteins listed in adjacent yellow table G – Green cluster consists of 72 proteins listed in green table below.

**Table 2:** Topological analysis of Overall protein-protein interaction network and the 4 clusters using the Cytohubba plugin.

|  | Rank | Node | Score | Rank | Node | Score | Rank | Node | Score |
| --- | --- | --- | --- | --- | --- | --- | --- | --- | --- |
| Overall | 1 | GRB2 | 64 | 1 | GRB2 | 118.83 | 1 | STAT3 | 3869.36 |
|  | 2 | **SRC** | 58 | 2 | SRC | 118 | 2 | EGFR | 3230.88 |
|  | 3 | PTPN11 | 54 | 3 | PTPN11 | 113.75 | 3 | FN1 | 3031.69 |
|  | 4 | SHC1 | 49 | 4 | EGFR | 113 | 4 | SRC | 2949.88 |
|  | 5 | EGFR | 48 | 5 | STAT3 | 109.5 | 5 | GRB2 | 2845.72 |
|  | 6 | STAT3 | 46 | 6 | SHC1 | 109.25 | 6 | KDR | 2137.02 |
|  | 7 | FYN | 42 | 7 | FYN | 105.92 | 7 | PTPN11 | 1981.46 |
|  | 8 | CBL | 41 | 8 | CBL | 105.83 | 8 | FYN | 1952.59 |
|  | 9 | FN1 | 40 | 9 | FN1 | 105.5 | 9 | CD44 | 1880.96 |
|  | 10 | **PLCG1** | 37 | 10 | KDR | 104.5 | 10 | LRP1 | 1867.42 |
| Red | 1 | GRB2 | 38 | 1 | GRB2 | 42.83 | 1 | GRB2 | 570.45 |
|  | 2 | PTPN11 | 30 | 2 | PTPN11 | 38.83 | 2 | FYN | 498.54 |
|  | 3 | SHC1 | 30 | 3 | SHC1 | 38.83 | 3 | PTPN11 | 186.99 |
| Blue | 1 | STAT3 | 28 | 1 | STAT3 | 32.12 | 1 | STAT3 | 573.05 |
|  | 2 | JAK1 | 17 | 2 | JAK1 | 26.62 | 2 | TRAF6 | 536.25 |
|  | 3 | **PDGFB** | 13 | 3 | SRC | 25.25 | 3 | SRC | 251.66 |
| Yellow | 1 | NOTCH1 | 11 | 1 | NOTCH1 | 13.83 | 1 | NOTCH1 | 158.33 |
|  | 2 | JAG1 | 7 | 2 | ARRB2 | 11.5 | 2 | ARRB2 | 73.2 |
|  | 3 | NOTCH2 | 7 | 3 | ARRB1 | 11 | 3 | ARRB1 | 42.8 |
| Green | 1 | FN1 | 26 | 1 | FN1 | 44.25 | 1 | FN1 | 1363.61 |
|  | 2 | ITGAM | 19 | 2 | ITGAM | 40.25 | 2 | ITGAM | 554.26 |
|  | 3 | **ITGB2** | 18 | 3 | CD44 | 38.67 | 3 | LRP1 | 549.81 |

### PPI network clustering identifies 4 functionally distinct groups of proteins

The k-means clustering algorithm identified 4 clusters (arbitrarily labelled red, blue, yellow and green; figure 3C to G) from within the overall PPI network. According to the topological analysis, GRB2, STAT3, NOTCH1 and FN1 were the top connected proteins in the red, blue, yellow and green clusters, respectively (table 2, figure S5). These clusters were significantly enriched for proteins related to ‘Protein kinase B signalling’ (red), ‘regulation of lymphocyte activation’ (blue), ‘Notch signalling’ (yellow) and ‘extracellular matrix organisation’ (green). In addition, KEGG pathway analysis identified ‘ErbB signalling’, ‘JAK-STAT signalling’, ‘Notch signalling’ and ‘coagulation and complement cascades’ pathways as significantly enriched in the red, blue, yellow and green clusters, respectively. All seed proteins that associated with AD belonged to the red (ErbB) cluster. Seed proteins associated with native T1 were mainly within red (ErbB) and yellow (Notch) PPI clusters and ECV within yellow (Notch) and green (extracellular matrix organisation) clusters. Proteins associated with LGE belonged to all four clusters (Table S2, column1).

## Discussion

Sensitive cardiovascular imaging has emerged as a key method to detect CV involvement in RA. We report on a first proteomic study of an early RA-CMR phenotyped cohort with time-linked vascular and myocardial CMR endpoints. Firstly, we identified 54 distinct proteins that were associated with CMR measures of VS and/or affected myocardial tissue in individuals with early, treatment naïve RA. Secondly, we discovered baseline proteins that were sensitive to change and associated with CMR changes over time, suggesting biological plausibility and opportunities in diagnostics and monitoring. Thirdly, PPI networks within this axis identified principal immune and coagulation pathways of ERA-CVD, including the JAK-STAT signalling pathway.

To our knowledge, this is the first longitudinal analysis of inflammation and cardio-vascular and -metabolic protein biomarkers to identify proteomic signatures of VS and affected myocardial tissue in a highly phenotyped, treatment naïve (no previous DMARD and/or corticosteroid use), active RA cohort with low CV risk (also excluding DM). CMR is a highly effective imaging modality, not least due to its multi-parametric capability, including tissue characterisation and quantitative markers of tissue inflammation, oedema and fibrosis^25^. This study’s parent CADERA trial reported VS (AD) and diffuse myocardial fibrosis (ECV measure) to be abnormal compared to controls at diagnosis ^9^. VS predicts major CV events independent of traditional CV risk score in people without CVD ^26^ ^27^. Similarly, myocardial fibrosis, a hallmark of HF, including with preserved ejection fraction (HFpEF) identified in people with RA^16^, is a poor prognostic marker in general populations ^28^ and can be detected on CMR prior to symptom onset ^29^. With inflammation a key contributor to increased risk of CVD in the RA population ^30^, we included an INF panel in addition to CV2, CV3 and CM Olink panels. No previous studies in RA have applied this breadth of proteins, evaluated proteins associated with VS as assessed on CMR or investigated an ERA cohort. Ahlers et al applied the Olink INF panel but in established and controlled RA and HF^16^ and identified proteins including ARTN associated with CMR-measured cardiac structure and function. In a general population cohort, there has been only one proteomic study (REMODEL) that identified differentially expressed proteins associated with measures of myocardial fibrosis (ECV and LGE) in patients with HTN and HTN+DM using the Olink CV2 and CV3 panels ^31^.

The 54 proteins identified at baseline in the current study included PDGF (platelet derived growth factor) subunit B and F11 (coagulation factor 11) that were associated with measures of VS; and ADAM-TS13 (a disintegrin and metalloproteinase with thrombospondin motifs 13), ARTN (Artemin) and PSGL-1 (P-selectin glycoprotein ligand-1) that associated with measures of myocardial fibrosis. Reduced expression of PDGF has been implicated in age-related endothelial dysfunction ^32^. A recent genome-wide association study (GWAS) from the UK Biobank also validated the association of PDGF signalling pathway with CMR-detected AD ^33^. A recent animal model study reported that the proteolytic activity of F11 needed for activation of extracellular matrix proteins in the heart inhibits genes involved in inflammation and fibrosis ^34^, suggesting a protective role of F11 in CVD. Upregulation of ADAM-TS13, a von Willebrand factor (vWF) protease, has been demonstrated previously in coronary plaques of individuals sustaining a myocardial infarction (MI) ^35^. In addition, in murine HF and ischaemic stroke models, administration of recombinant human and constitutively active variant of ADAM-TS13 have been shown separately to reduce inflammation, improve myocardial remodelling and function^36^, and significantly improve regional cerebral blood flow and reduce lesion volume respectively ^37^. ARTN, expressed by vascular (including coronary) smooth muscle cells ^38^, has been shown to be differentially expressed in RA versus healthy individuals and positively associated with CMR measured ventricular-vascular coupling ratio ^16^ that in turn is associated with HF development ^39^. PSGL-1, a glycoprotein expressed on all leucocytes including monocytes, is critical for the formation of monocyte-platelet complexes (MPCs) that have been correlated with severity and prognosis of CVD^40,41^ and associated with increased high-sensitivity CRP (hsCRP)^42^. These proteins provide valuable insights into potential biomarkers for CVD risk stratification and early detection in (early) RA and therapeutic targets.

The study also identified proteins TNFSF13B [also known as B-cell activating factor (BAFF)] and CRTAC1 that were sensitive to change over time, which importantly, associated with interval change in CMR measures. The role of BAFF in CVD has not yet been fully elucidated. In preclinical models of atherosclerosis, B-cell depletion has been shown to reduce atherogenesis and improve cardiac recovery post MI^43^ whilst anti-BAFF antibodies have been demonstrated as pro-atherogenic^44^. The CADERA trial^9^ reported AD improved longitudinally in response to treatment. In the current study, BAFF was associated with AD over time and thus may be regarded as a potential biomarker for AD. CRTAC1, on the other hand, was positively associated with ECV change. ECV is a measure of interstitial myocardial fibrosis and a marker of myocardial tissue remodelling. No studies on CVD have implicated this protein thus far. However, it has been reported as a candidate biomarker for osteoarthritis and a predictor of progression to joint replacement ^45^. Interestingly, only 2 inflammation-related proteins SIRT2 and STAMBP were associated with myocardial fibrosis (ECV) at baseline. By year 1, this association changed significantly to a negative one suggestive of their dynamic role in disease pathogenesis modulated either through the effects of treatment or evolving disease activity.

Proteins involved in EGFR tyrosine kinase inhibitor resistance and JAK-STAT signalling were significantly enriched in the expanded protein network. The ORAL-Surveillance study (OSS) ^46^ demonstrated tofacitinib (inhibitor of JAK1, 3) was non-inferior to a TNF inhibitor in established RA patients (age > 50 years + 1 additional CV risk factor). The basis for this and the role of JAK isoforms is unclear although some studies implicate the JAK2 isoform in atherogenesis^47,48^. The current study reported proteins involved in JAK-STAT signalling within the overall ERA-CVD network (STAT3) and the blue PPI cluster (JAK1 and STAT3). In addition, Growth factor receptor bound protein 2 (GRB2) protein was the most connected within the network and is involved in the two top signalling pathways – EGFR tyrosine kinase and JAK-STAT signalling. While it has not been directly implicated in CVD, a related protein, GRB2 associated binding proteins-2 (GAB-2) has been reported recently as a key regulator of inflammatory signalling that leads to vascular inflammation and thrombosis ^49^. Additional clusters of proteins were identified, involved in key processes such as ErbB (red cluster) and Notch signalling (yellow cluster) that were associated with measures of VS and myocardial tissue respectively.

It is important to acknowledge the limitations of our study. Pre-defined Olink panels of proteins relevant to inflammation and cardiovascular diseases were applied rather than comprehensive sampling of the entire proteome. To partly circumvent this, the seed ERA-CVD PPI network was expanded to identify other proteins of relevance to this axis. Alternative approaches, such as Olink Explore, SomaScan^50^ and shotgun proteomic techniques ^51^, offer broader proteomic coverage and could unveil additional protein associations that our study might have missed. In focusing on the primary and key secondary CMR pathophysiological endpoints of the CADERA study, this analysis did not capture all CMR measures such as those investigating the myocardial structure and function. Other CV imaging modalities offer evaluation of other aspects of CV pathophysiology that would be useful to also investigate. Validating our findings in an independent cohort would be a next step to support generalizability of our results and inform clinical translation.

In summary, this study identifies new protein biomarkers associated with subclinical CMR-defined abnormalities in VS and myocardial tissue and sensitivity to change, in an ERA cohort with low CV risk profile. Key signalling pathways of the ERA-CVD multimorbid axis, including JAK-STAT signalling are identified. Validating these findings would provide a basis for future risk stratification and therapeutic target discovery.

## Author Contributions

RRS and MHB had full access to all the data in the study and takes responsibility for the integrity of the data and the accuracy of the data analysis.

*Concept and design: RRS, DP, MHB*

*Acquisition, analysis, or interpretation of data: All authors.*

*Drafting of the manuscript: RRS drafted the manuscript with critical input from DP and MHB.*

*All authors had the opportunity to further revise the manuscript and approved the final version.*

*Critical review of the manuscript for important intellectual content: MHB, DP, MHB*

*Statistical analysis: RRS, DP, MHB.*

*Obtained funding: MHB.*

*Administrative, technical, or material support: All authors.*

*Supervision: DP, MHB.*

## Data Sharing Statement

*Data are available upon reasonable request. All data relevant to the study are included in the article or uploaded as supplementary information. Additional data are available upon reasonable request.*

## Supplementary material

Table S1: Proteins below the lower limit of detection (LLOD) for each Olink panel

Table S2: Baseline Protein NPX (normalised protein expression) posterior estimates with 95% credible intervals.

Table S3: Proteins sensitive to change over time.

Table S4: Protein NPX*time interaction posterior estimates with 95% credible intervals.

Table S5: Structural properties of the expanded eRA-CVD PPI network

Figure S1: Correlation matrix of 54 proteins (raw data) and clinically measured high sensitivity troponin I (HsTnI_CLIN) and NTProBNP (NTProBNP_CLIN).

Figure S2: Top = Predicted NPX for CRTAC1 (left) and TNFSF13B at baseline and year 1 Bottom = Partial residuals plot for interaction between NPX*time for CRTAC1 and myocardial extracellular volume; TNFSF13B and aortic distensibility.

Figure S3: Partial residuals plot for interaction between NPX*time for SIRT2 (top) and STAMBP (bottom) and myocardial extracellular volume.

Figure S4: Top – Top 10 GO Biological processes (left) and KEGG pathways (right)

Figure S5: Top 10 GO Biological and KEGG pathways for 4 protein-protein interaction (PPI) network clusters

---

**Table S1:**
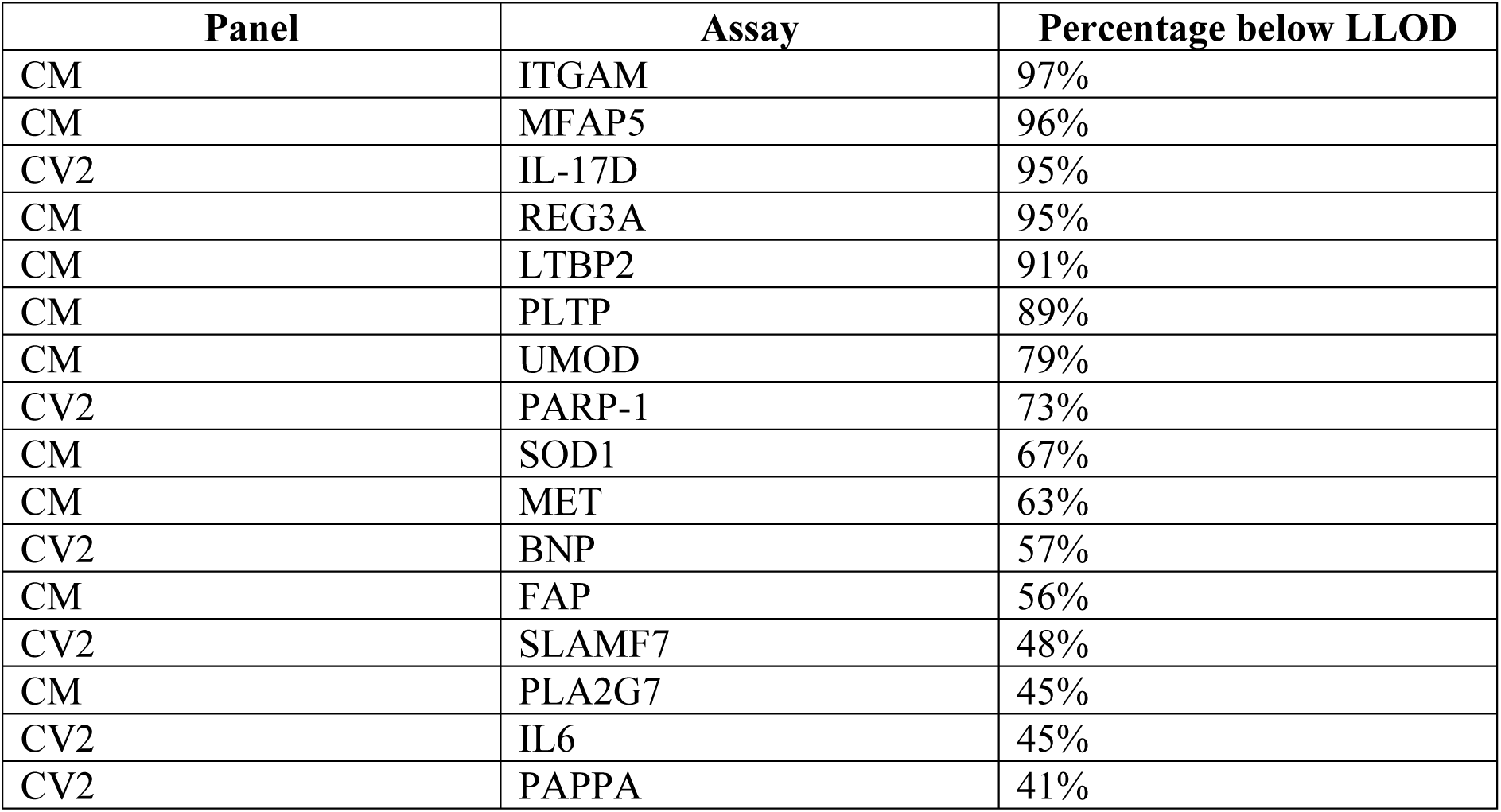
Proteins below the lower limit of detection (LLOD) for each Olink panel.

| Panel | Assay | Percentage below LLOD |
| --- | --- | --- |
| CM | ITGAM | 97% |
| CM | MFAP5 | 96% |
| CV2 | IL-17D | 95% |
| CM | REG3A | 95% |
| CM | LTBP2 | 91% |
| CM | PLTP | 89% |
| CM | UMOD | 79% |
| CV2 | PARP-1 | 73% |
| CM | SOD1 | 67% |
| CM | MET | 63% |
| CV2 | BNP | 57% |
| CM | FAP | 56% |
| CV2 | SLAMF7 | 48% |
| CM | PLA2G7 | 45% |
| CV2 | IL6 | 45% |
| CV2 | PAPPA | 41% |

**Table S2:** Baseline Protein NPX posterior estimates with 95% credible intervals. In descending order of strength of association for each CMR measure. ECV = extracellular volume, LGE – late gadolinium enhancement, AD = aortic distensibility, AoSI = aortic stiffness index. CM = cardiometabolic panel, CV2 = cardiovascular II panel, CV3 = cardiovascular III panel, INF = inflammation panel.

| PPI cluster | CMR measure | Assay | Panel | Posterior estimate | 95%credible interval (upper limit) | 95%credible interval (lower limit) | % imputed |
| --- | --- | --- | --- | --- | --- | --- | --- |
| Green | Native T1 | ADAM-TS13 | CV2 | -85.05 | -11.41 | -158.69 | 0 |
| Red | Native T1 | PLC | CV3 | -39.35 | -1.38 | -77.33 | 1 |
| Red | Native T1 | GDNF | INF | 35.38 | 69.76 | 1.01 | 1 |
| Yellow | Native T1 | TR-AP | CV3 | -35.24 | -5.83 | -64.64 | 1 |
| Yellow | Native T1 | IDUA | CV2 | -28.92 | -3.83 | -54.01 | 0 |
| Yellow | Native T1 | RAGE | CV2 | -27.75 | -2.02 | -53.48 | 0 |
| Yellow | Native T1 | ST2 | CV3 | -24.43 | -6.23 | -42.62 | 1 |
| Red | Native T1 | CD5 | INF | 24.37 | 47.63 | 1.12 | 0 |
| Blue | Native T1 | TNFRSF9 | INF | 19.41 | 38.58 | 0.24 | 0 |
| Red | ECV | ARTN | INF | -0.0393 | -0.0083 | -0.0704 | 34 |
| Red | ECV | NRP1 | CM | 0.0349 | 0.0597 | 0.01 | 0 |
| Green | ECV | CTSD | CV3 | -0.0318 | -0.0131 | -0.0506 | 1 |
| Green | ECV | F11 | CM | -0.0287 | -0.0039 | -0.0535 | 0 |
| Green | ECV | COL18A1 | CM | -0.0272 | -0.0057 | -0.0487 | 0 |
| Green | ECV | SERPINA5 | CM | -0.027 | -0.01 | -0.044 | 0 |
| Green | ECV | IGFBP3 | CM | -0.0246 | -0.0063 | -0.0428 | 0 |
| Yellow | ECV | CRTAC1 | CM | 0.0231 | 0.0369 | 0.0093 | 0 |
| Green | ECV | MMP-2 | CV3 | 0.0216 | 0.0432 | 0.0001 | 1 |
| Green | ECV | PCOLCE | CM | -0.0213 | -0.0031 | -0.0395 | 0 |
| Green | ECV | LDL receptor | CV3 | -0.0203 | -0.0069 | -0.0338 | 1 |
| Yellow | ECV | PRSS8 | CV2 | -0.0197 | -0.0041 | -0.0353 | 0 |
| Green | ECV | FCN2 | CM | -0.0188 | -0.0048 | -0.0329 | 0 |
| Yellow | ECV | Notch 3 | CV3 | 0.0186 | 0.0361 | 0.0012 | 1 |
| Green | ECV | t-PA | CV3 | -0.0181 | -0.0073 | -0.0289 | 1 |
| Blue | ECV | TNFSF13B | CV3 | 0.0169 | 0.0325 | 0.0013 | 1 |
| Yellow | ECV | TIMD4 | CM | 0.0168 | 0.0306 | 0.0031 | 0 |
| Green | ECV | ANG | CM | -0.0166 | -0.0023 | -0.0309 | 0 |
| Yellow | ECV | CNDP1 | CM | -0.0162 | -0.0005 | -0.0318 | 0 |
| Blue | ECV | IL7R | CM | 0.0135 | 0.027 | 0.005 | 4 |
| Blue | ECV | IL-20RA | INF | -0.0118 | -0.0002 | -0.0234 | 21 |

Table S2: Baseline Protein NPX posterior estimates with 95% credible intervals. In descending order of strength of association for each CMR measure. ECV = extracellular
| PPI cluster (continued) | CMR measure (continued) | Assay (continued) | Panel (continued) | Posterior estimate (continued) | 95%credible interval (upper limit) (continued) | 95%credible interval (lower limit) (continued) | % imputed (continued) |
| --- | --- | --- | --- | --- | --- | --- | --- |
| Yellow | ECV | FABP4 | CV3 | -0.0111 | -0.0013 | -0.0209 | 1 |
| Green | ECV | CA5A | CV2 | -0.011 | -0.0015 | -0.0219 | 14 |
| Green | ECV | TGM2 | CV2 | 0.0109 | 0.0183 | 0.0036 | 0 |
| NA | ECV | NT-proBNP | CV3 | 0.0106 | 0.0204 | 0.0008 | 6 |
| Red | ECV | STAMBP | INF | 0.0095 | 0.0171 | 0.0019 | 0 |
| Red | ECV | FGF-19 | INF | 0.0084 | 0.0168 | 0.0001 | 0 |
| Yellow | ECV | SIRT2 | INF | 0.0071 | 0.013 | 0.0012 | 0 |
| Green | ECV | MMP-3 | CV3 | 0.0063 | 0.0122 | 0.0004 | 1 |
| Red | ECV | HAOX1 | CV2 | -0.0043 | -0.0003 | -0.0084 | 0 |
| Green | LGE | PSGL-1 | CV2 | -2.73 | -0.05 | -5.4 | 0 |
| Green | LGE | ICAM-2 | CV3 | -2.71 | -0.18 | -5.24 | 1 |
| Yellow | LGE | ITGB1BP2 | CV2 | 2.09 | 3.83 | 0.35 | 40 |
| Blue | LGE | IL-12B | INF | -1.76 | -0.34 | -3.18 | 0 |
| Blue | LGE | SRC | CV2 | 1.54 | 2.76 | 0.31 | 0 |
| Red | LGE | STK4 | CV2 | 1.34 | 2.55 | 0.13 | 2 |
| Red | LGE | GT | CV2 | 1.29 | 2.51 | 0.08 | 1 |
| Yellow | LGE | CCL3 | CV2 | 1.17 | 2.17 | 0.16 | 0 |
| Red | LGE | FGF-21 | INF | -0.93 | 0 | -1.86 | 0 |
| Red | AD | PDGF subunit B | CV2 | -2.65 | -0.36 | -4.94 | 0 |
| Red | AD | FGF-23 | CV2 | 0.96 | 1.89 | 0.03 | 5 |
| Red | AD | HB-EGF | CV2 | -0.6 | -0.05 | -1.14 | 0 |
| Green | AoSI | F11 | CM | -1.8 | -0.07 | -3.53 | 0 |
| Green | AoSI | PCSK9 | CV3 | -1.65 | -0.33 | -2.98 | 1 |
| Yellow | AoSI | PI3 | CV3 | 1.12 | 2.1 | 0.15 | 1 |
| Red | AoSI | HB-EGF | CV2 | 0.9 | 1.63 | 0.18 | 0 |
| Yellow | AoSI | SCGB3A2 | CV3 | -0.57 | -0.11 | -1.03 | 1 |

**Table S3:** Proteins sensitive to change over time. *Proteins also associated with CMR measures at baseline ** Proteins also associated with CMR measures at baseline and with a significant interaction between baseline-year1 NPX. CM = cardiometabolic panel, CV2 = cardiovascular II panel, CV3 = cardiovascular III panel, INF = inflammation panel.

| Assay | Panel | Posterior estimate | 95%credible interval (upper limit) | 95%credible interval (lower limit) |
| --- | --- | --- | --- | --- |
| TNF-R2 | CV3 | 4.378 | 5.09 | 3.665 |
| **MMP-3\*** | **CV3** | **-1.08** | **-0.818** | **-1.343** |
| **CCL3\*** | **CV2** | **-0.554** | **-0.395** | **-0.713** |
| PRTN3 | CV3 | -0.554 | -0.357 | -0.75 |
| IL16 | CV2 | -0.515 | -0.373 | -0.658 |
| AZU1 | CV3 | -0.513 | -0.268 | -0.758 |
| SAA4 | CM | -0.455 | -0.339 | -0.571 |
| IL2-RA | CV3 | -0.442 | -0.335 | -0.549 |
| CCL18 | CM | -0.414 | -0.245 | -0.582 |
| CFHR5 | CM | -0.411 | -0.258 | -0.564 |
| CHI3L1 | CV3 | -0.403 | -0.218 | -0.588 |
| CEACAM8 | CV2 | -0.402 | -0.218 | -0.585 |
| IL-1ra | CV2 | -0.386 | -0.235 | -0.537 |
| IGLC2 | CM | -0.364 | -0.278 | -0.45 |
| MMP-9 | CV3 | -0.349 | -0.215 | -0.483 |
| MMP12 | CV2 | -0.341 | -0.227 | -0.455 |
| PGLYRP1 | CV3 | -0.324 | -0.21 | -0.438 |
| LOX-1 | CV2 | -0.321 | -0.168 | -0.475 |
| CXCL1 | CV2 | -0.321 | -0.194 | -0.447 |
| IL-10RA | INF | -0.317 | -0.03 | -0.605 |
| MPO | CV3 | -0.31 | -0.173 | -0.448 |
| **HB-EGF\*** | **CV2** | **-0.307** | **-0.193** | **-0.422** |
| CDCP1 | INF | 0.301 | 0.527 | 0.076 |
| SELE | CV3 | -0.299 | -0.211 | -0.388 |
| THBS4 | CM | 0.299 | 0.583 | 0.014 |
| **TIMD4\*** | **CM** | **-0.284** | **-0.188** | **-0.379** |
| RETN | CV3 | -0.281 | -0.173 | -0.389 |
| **FCN2\*** | **CM** | **-0.28** | **-0.214** | **-0.346** |
| ITGB2 | CV3 | -0.272 | -0.169 | -0.375 |
| DPP4 | CM | 0.269 | 0.355 | 0.183 |
| PDGF subunit A | CV3 | -0.259 | -0.167 | -0.352 |
| IGFBP-2 | CV3 | -0.259 | -0.112 | -0.406 |
| U-PAR | CV3 | -0.252 | -0.156 | -0.348 |
| CXCL5 | INF | 0.248 | 0.473 | 0.022 |
| TR | CV3 | -0.247 | -0.049 | -0.445 |
| TNF-R1 | CV3 | -0.239 | -0.147 | -0.331 |
| CD163 | CV3 | -0.238 | -0.141 | -0.334 |
| **SERPINA5\*** | **CM** | **0.211** | **0.31** | **0.113** |
| TLT-2 | CV3 | -0.21 | -0.107 | -0.312 |
| ST6GAL1 | CM | -0.207 | -0.087 | -0.328 |
| LCN2 | CM | -0.205 | -0.083 | -0.326 |
| GRN | CV3 | -0.202 | -0.139 | -0.265 |
| ICAM1 | CM | -0.199 | -0.123 | -0.275 |
| MB | CV3 | 0.196 | 0.355 | 0.037 |
| **CRTAC1\*\*** | **CM** | **-0.192** | **-0.104** | **-0.28** |
| FS | CV2 | -0.189 | -0.062 | -0.316 |
| RARRES2 | CV3 | -0.188 | -0.129 | -0.247 |
| AOC3 | CM | 0.187 | 0.253 | 0.12 |
| CTSL1 | CV2 | -0.185 | -0.085 | -0.284 |
| TNFRSF13B | CV2 | -0.184 | -0.114 | -0.255 |
| C2 | CM | -0.184 | -0.124 | -0.244 |
| LPL | CV2 | 0.184 | 0.276 | 0.092 |
| GDF-2 | CV2 | 0.181 | 0.308 | 0.054 |
| IL-18BP | CV3 | -0.179 | -0.091 | -0.266 |

Table S3: Proteins sensitive to change over time.
| Assay (continued) | Panel (continued) | Posterior estimate (continued) | 95%credible interval (upper limit) (continued) | 95%credible interval (lower limit) (continued) |
| --- | --- | --- | --- | --- |
| NCAM1 | CM | 0.176 | 0.252 | 0.1 |
| PAI | CV3 | -0.173 | -0.093 | -0.253 |
| Gal-3 | CV3 | -0.172 | -0.108 | -0.236 |
| TIMP1 | CM | -0.17 | -0.099 | -0.24 |
| CNTN1 | CV3 | 0.164 | 0.244 | 0.085 |
| GNLY | CM | -0.163 | -0.055 | -0.271 |
| THPO | CV2 | -0.16 | -0.073 | -0.247 |
| C1QTNF1 | CM | 0.16 | 0.272 | 0.048 |
| **IDUA\*** | **CV2** | **0.156** | **0.241** | **0.071** |
| CD4 | CV2 | -0.154 | -0.096 | -0.213 |
| **CNDP1\*** | **CM** | **0.15** | **0.227** | **0.072** |
| ANGPLT3 | CM | -0.148 | -0.044 | -0.251 |
| **NRP1\*** | **CM** | **-0.144** | **-0.062** | **-0.227** |
| TFPI | CV3 | -0.141 | -0.079 | -0.204 |
| TF | CV2 | 0.139 | 0.197 | 0.081 |
| **TNFSF13B\*\*** | **CV3** | **-0.137** | **-0.047** | **-0.227** |
| TNFRSF11A | CV2 | -0.137 | -0.054 | -0.219 |
| SCF | CV2 | 0.131 | 0.217 | 0.045 |
| MARCO | CV2 | -0.129 | -0.079 | -0.179 |
| TNFRSF14 | CV3 | -0.127 | -0.038 | -0.217 |
| **TR-AP\*** | **CV3** | **-0.119** | **-0.053** | **-0.185** |
| TRAIL-R2 | CV2 | -0.113 | -0.035 | -0.191 |
| Dkk-1 | CV2 | -0.112 | -0.037 | -0.187 |
| IL-4RA | CV2 | -0.101 | -0.031 | -0.172 |
| LTBR | CV3 | -0.098 | -0.028 | -0.168 |
| OSMR | CM | -0.093 | -0.049 | -0.137 |
| VCAM1 | CM | -0.083 | -0.028 | -0.138 |
| hOSCAR | CV2 | -0.077 | -0.027 | -0.128 |
| SPON1 | CV3 | -0.075 | -0.022 | -0.129 |
| AMBP | CV2 | -0.067 | -0.035 | -0.099 |
| PRELP | CV2 | 0.06 | 0.103 | 0.018 |

**Table S4:** Protein NPX*time interaction posterior estimates with 95% credible intervals. In descending order of strength of association. *Proteins also associated with CMR measures at baseline ** Proteins also associated with CMR measures at baseline and sensitive to change over time ECV = extracellular volume, LGE – late gadolinium enhancement, AD = aortic distensibility, AoSI = aortic stiffness index. CM = cardiometabolic panel, CV2 = cardiovascular II panel, CV3 = cardiovascular III panel, INF = inflammation panel.

| CMR measure | Assay | Panel | Posterior estimate for NPX*week | 95%credible interval (upper limit) | 95%credible interval (lower limit) | % imputed |
| --- | --- | --- | --- | --- | --- | --- |
| Native T1 | IL-15RA | INF | -64.27 | -18.60 | -109.93 | 1 |
| Native T1 | AOC3 | CM | 47.06 | 92.53 | 1.59 | 0 |
| Native T1 | NT-3 | INF | -31.69 | -1.27 | -62.11 | 0 |
| Native T1 | CCL25 | INF | -24.85 | -1.28 | -48.42 | 0 |
| **Native T1** | **STAMBP\*** | **INF** | **-23.20** | **-4.68** | **-41.72** | **0** |
| **Native T1** | **SIRT2\*** | **INF** | **-15.41** | **-1.12** | **-29.69** | **0** |
| **ECV** | **CRTAC1\*\*** | **CM** | **-0.0179** | **-0.0021** | **-0.0338** | **0** |
| ECV | CCL5 | CM | 0.0152 | 0.0276 | 0.0027 | 0 |
| ECV | CCL4 | INF | -0.0151 | -0.0041 | -0.0262 | 0 |
| **ECV** | **STAMBP\*** | **INF** | **-0.0131** | **-0.0013** | **-0.0248** | **0** |
| **ECV** | **SIRT2\*** | **INF** | **-0.0091** | **-0.0007** | **-0.0174** | **0** |
| ECV | SERPINA12 | CV2 | 0.0074 | 0.0142 | 0.0005 | 0 |
| AD | PAM | CM | -2.16 | -0.36 | -3.96 | 0 |
| **AD** | **COL18A1\*** | **CM** | **-1.8** | **-0.5** | **-3.11** | **0** |
| AD | CCL14 | CM | -1.76 | -0.53 | -2.99 | 0 |
| AD | LIF-R | INF | -1.75 | -0.02 | -3.47 | 0 |
| AD | ST6GAL1 | CM | -1.58 | -0.34 | -2.82 | 0 |
| AD | CST5 | INF | -1.52 | -0.69 | -2.35 | 0 |
| **AD** | **PCOLCE\*** | **CM** | **-1.45** | **-0.4** | **-2.5** | **0** |
| AD | TIMP1 | CM | -1.45 | -0.15 | -2.74 | 0 |
| **AD** | **FGF-23\*** | **CV2** | **1.41** | **2.58** | **0.24** | **5** |
| AD | PD-L2 | CV2 | -1.36 | -0.07 | -2.66 | 1 |
| AD | MCP-1 | CV3 | -1.24 | -0.35 | -2.14 | 1 |
| AD | CST3 | CM | -1.21 | -0.14 | -2.28 | 0 |
| AD | Flt3L | INF | -1.15 | -0.21 | -2.1 | 0 |
| **AD** | **TNFSF13B\*\*** | **CV3** | **-1.09** | **-0.05** | **-2.18** | **1** |
| AD | TIMP4 | CV3 | 1.07 | 1.81 | 0.32 | 1 |
| AD | ANGPTL3 | CM | -1.04 | -0.11 | -1.98 | 0 |
| AD | CXCL1 | CV2 | 0.92 | 1.58 | 0.25 | 0 |
| AD | IGFBP-2 | CV3 | -0.73 | -0.2 | -1.25 | 1 |
| AD | CCL18 | CM | -0.58 | -0.06 | -1.1 | 0 |

Table S4: Protein NPX\*time interaction posterior estimates with 95% credible intervals. In descending order of strength of association.
| CMR measure (continued) | Assay (continued) | Panel (continued) | Posterior estimate for NPX*week (continued) | 95%credible interval (upper limit) (continued) | 95%credible interval (lower limit) (continued) | % imputed (continued) |
| --- | --- | --- | --- | --- | --- | --- |
| AoSI | PD-L2 | CV2 | 1.86 | 3.5 | 0.22 | 1 |
| **AoSI** | **IGFBP3\*** | **CM** | **1.58** | **2.99** | **0.18** | **0** |
| AoSI | PON3 | CV3 | -0.97 | -0.07 | -1.86 | 1 |

**Table S5:** Structural properties of the expanded early rheumatoid arthritis-cardiovascular disease protein-protein interaction network.

|  |  |
| --- | --- |
| No. of nodes | 203 |
| No. of edges | 1108 |
| Avg. no. of neighbours | 11.78 |
| Network diameter | 6 |
| Network radius | 3 |
| Characteristic path length | 2.78 |
| Clustering coefficient | 0.49 |
| Network density | 0.06 |
| Network heterogeneity | 0.98 |
| Network centralisation | 0.28 |
| Connected components | 15 |

**Figure S1:**
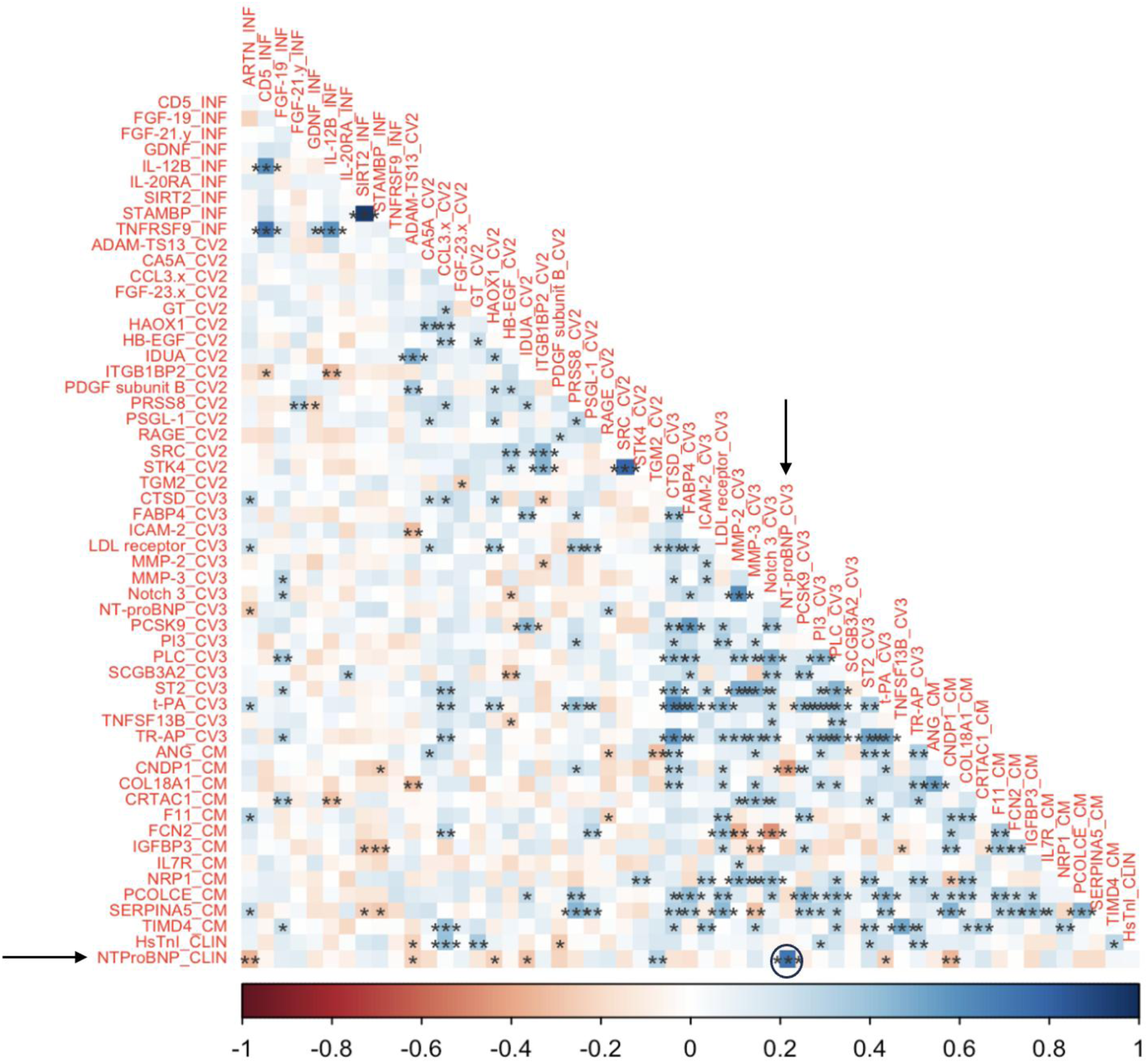
Correlation matrix of 54 proteins (raw data) and clinically measured high sensitivity troponin I (HsTnI_CLIN) and NTProBNP (NTProBNP_CLIN). Arrows draw attention to high correlation (coefficient = 0.77) of Olink and clinically measured NTProBNP with high significance. * p < 0.05, **p <0.01, ***p<0.005. CM = cardiometabolic panel, CV2 = cardiovascular II panel, CV3 = cardiovascular III panel, INF = inflammation panel.

**Figure S2:**
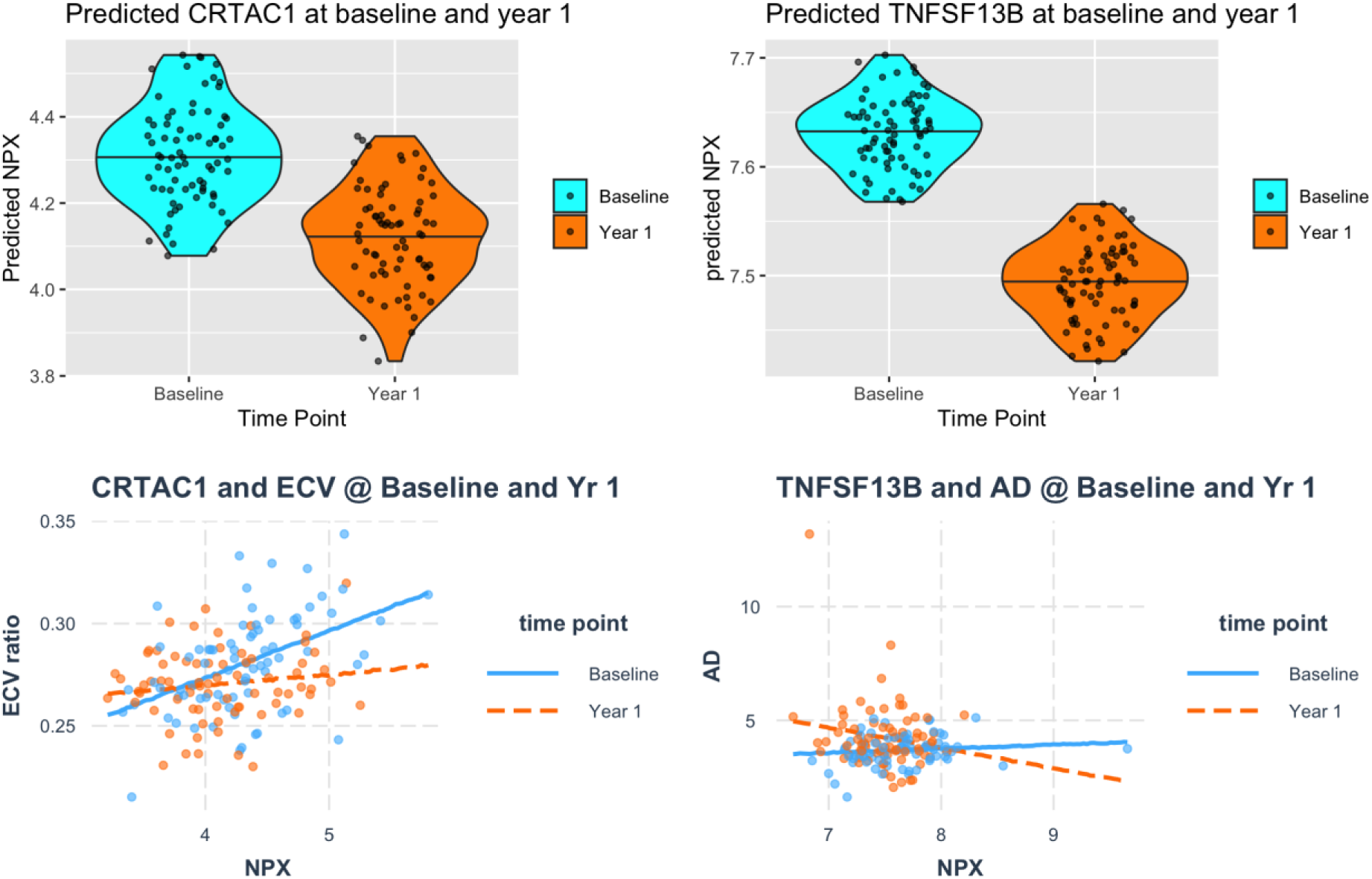
TOP = Predicted NPX for CRTAC1 (left) and TNFSF13B at baseline and year 1 (adjusted for age, gender, pack years smoked and systolic blood pressure). Bottom = Partial residuals plot for interaction between NPX*time for CRTAC1 and myocardial extracellular volume (left) and TNFSF13B and aortic distensibility (right). ECV = extracellular volume, AD = aortic distensibility.

**Figure S3:**
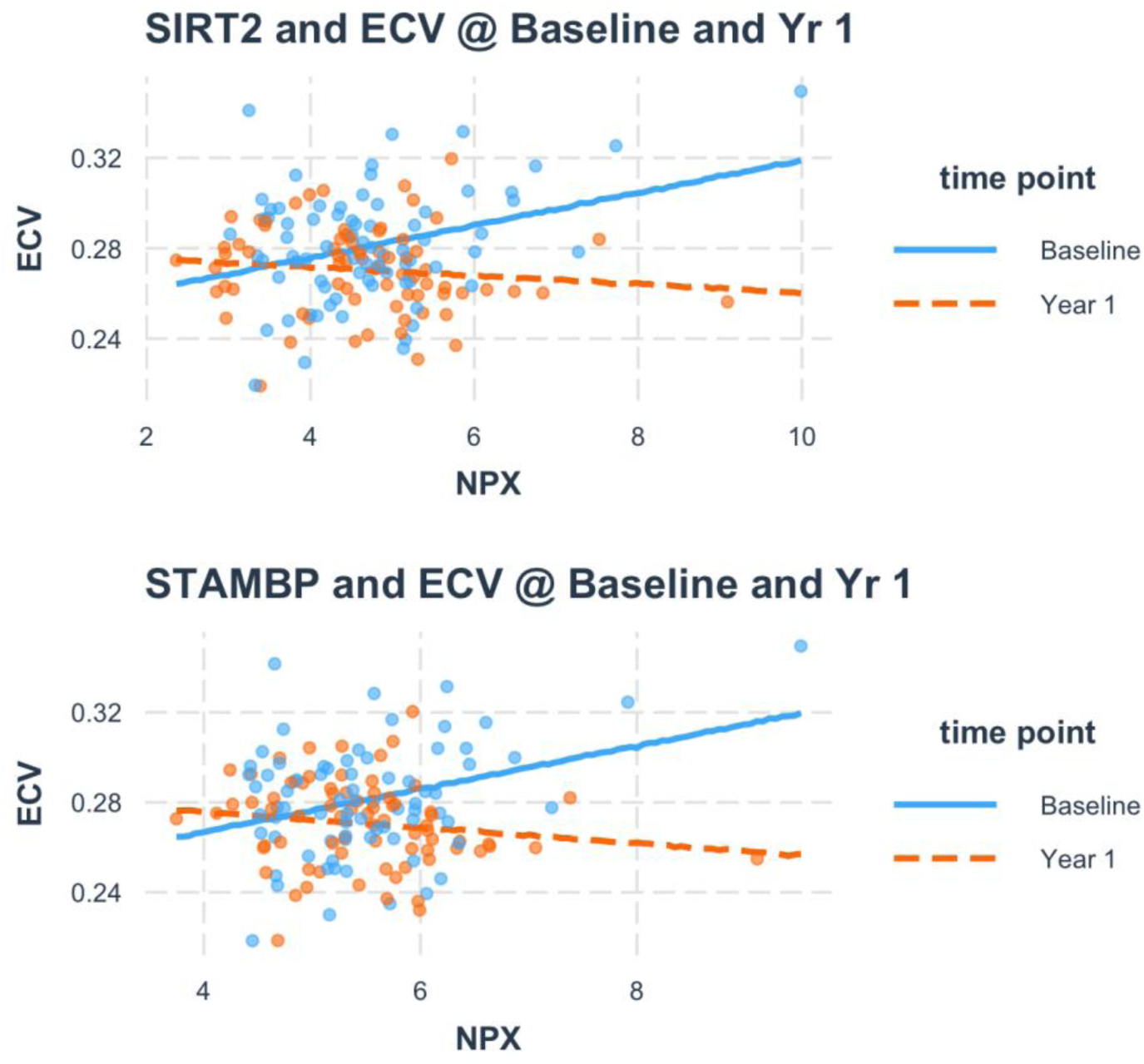
Partial residuals plot for interaction between NPX*time for SIRT2 (top) and STAMBP (bottom) and myocardial extracellular volume (ECV).

**Figure S4:**
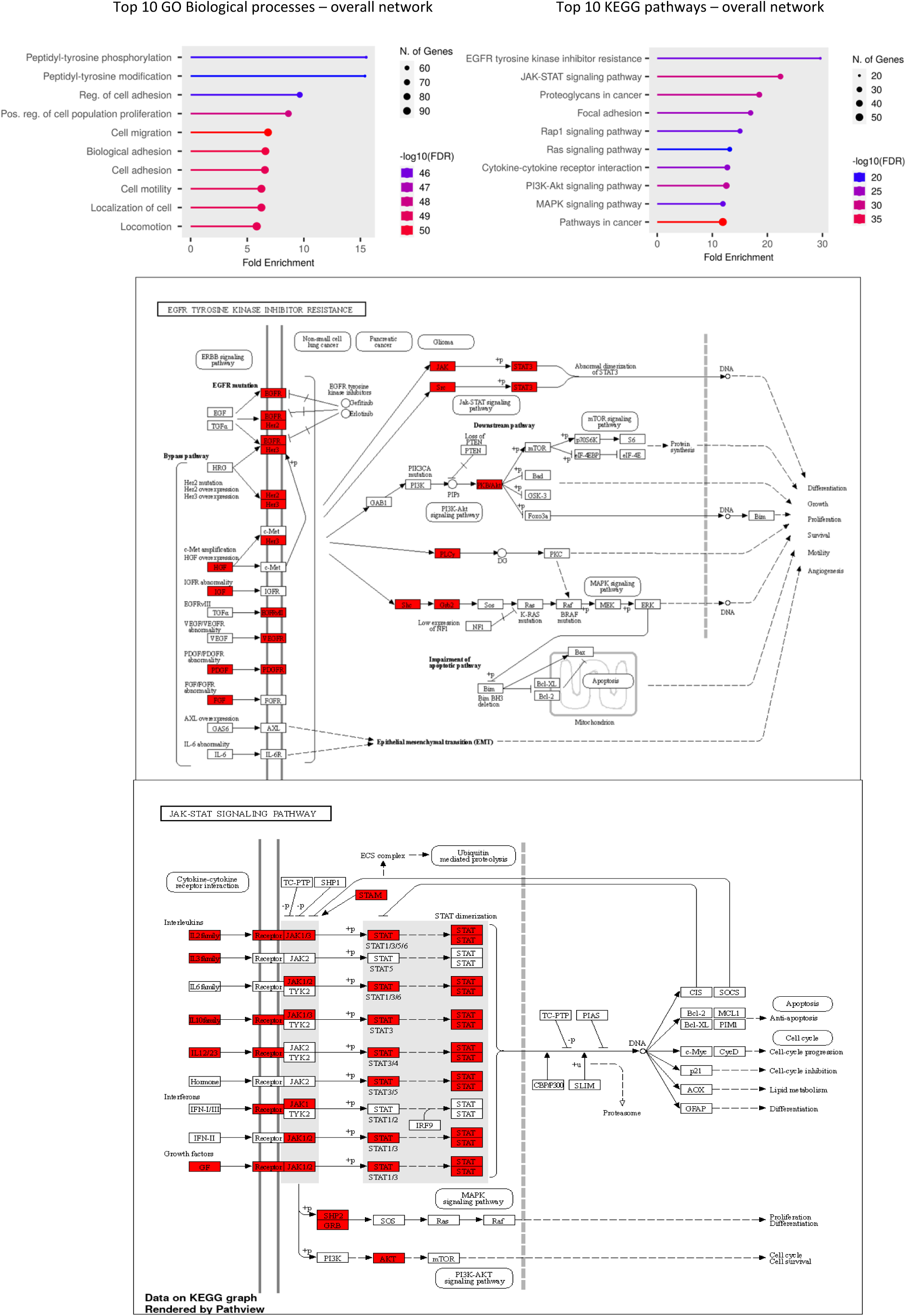
Top – Top 10 GO Biological processes (left) and KEGG pathways (right) Middle – EGFR tyrosine kinase inhibitor resistance pathway with proteins identified in overall network highlighted in Red. Bottom – JAK STAT signalling pathway with proteins identified in overall network highlighted in Red (Image generated using ShinyGo)

**Figure S5:**
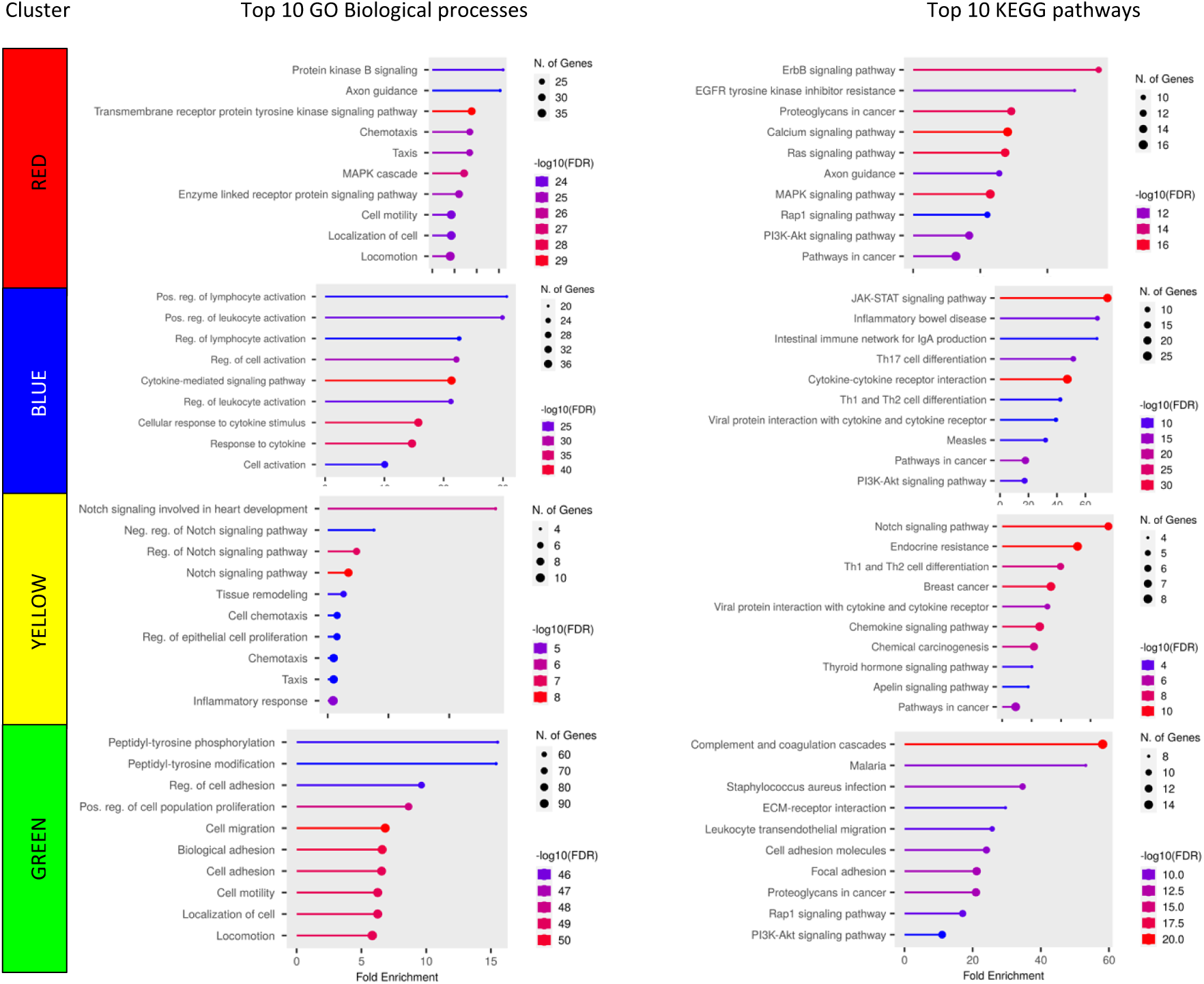
Top 10 GO Biological and KEGG pathways for 4 protein-protein interaction (PPI) network clusters. Fold enrichment on X axis, colour indicates −log10(FDR) and size of dot represents number of proteins.

